# Cytoophidia influence cell cycle and size in *Schizosaccharomyces pombe*

**DOI:** 10.1101/2023.10.15.562340

**Authors:** Ruolan Deng, Yi-Lan Li, Ji-Long Liu

## Abstract

CTP synthase (CTPS) forms cytoophidia in all three domains of life. Here we focus on the function of cytoophidia in cell proliferation using *Schizosaccharomyces pombe* as a model system. We find that converting His^359^ of CTPS into Ala^359^ leads to cytoophidium disassembly. By reducing the level of CTPS protein or specific point mutations, the loss of cytoophidia prolongs the G2 phase and expands cell size. In addition, the loss-filament mutant of CTPS leads to a decrease in the expression of genes related to G2/M transition and cell growth, including *slm9*. The overexpression of *slm9* alleviates the G2 phase elongation and cell size enlargement induced by CTPS loss-filament mutant. Overall, our results connect cytoophidia with cell cycle and cell size control in *Schizosaccharomyces pombe*.

## 1. Introduction

The nucleotide cytidine triphosphate (CTP) plays a crucial role in various biological processes, including RNA synthesis, phospholipid synthesis, protein glycosylation, and energy transfer enzymatic reactions across diverse organisms. CTPS is the rate-limiting enzyme to catalyzes the ATP-dependent conversion of UTP to CTP [1]. Notably, CTPS has been observed to form filamentous structures termed cytoophidia in a wide range of organisms from prokaryotes to eukaryotes. Cytoophidia have been identified in *Drosophila*, *Caulobacter crescentus*, and *S. cerevisiae* since 2010 [2–4], with subsequent studies confirming their presence in human cells [5][6], fission yeast [7], plants [8], archaea [9], and mammalian cells [10]. These findings highlight the highly conserved nature of cytoophidia among different organisms and suggest that they possess significant yet undiscovered biological functions.

The role of cytoophidia remains enigmatic, despite proposed physiological functions in various organisms. Multiple studies have demonstrated that cytoophidia regulate cell metabolism by adopting different conformations to control CTPS enzymatic activity across different organisms [11–15]. In *C. crescentus*, cytoophidia cooperated with the intermediate filament to maintain cellular shape [4]. In *Drosophila*, cytoophidia are involved in optic lobe development and oogenesis; recent studies showed that cytoophidia take part in adipose architecture and metabolism, and loss of cytoophidia reduced adipocyte expansion and inhibition of lipogenesis [15–18]. In human cells, cytoophidia stabilize the CTPS protein by extending its half-life [19]. Cytoophidia participate in ovarian stress response in *Drosophila* [20], as well as respond to changes in nutrition, pH, temperature, and osmolality for stress adaptation in yeast [21,22]. Furthermore, cytoophidia contributed to rapid cell proliferation, they have been observed in neuroblasts of *Drosophila* larvae [13] and highly proliferated T cells within the murine thymus [10]. Additionally, elevated CTPS activity has been found in various cancers such as hepatoma and lymphoma; moreover, cytoophidia have been identified within human cancers like hepatocellular carcinoma [5,23,24], suggesting an important role for cytoophidia in cell proliferation.

The cytoophidia are believed to play a crucial role in coupling cell proliferation, particularly in fast-growing cells like cancer cells and activated T cells [5,10]. Notably, studies have shown that mouse embryonic stem cells possess abundant cytoophidia [25]. In the context of cancer, the formation of cytoophidia may indicate an abnormal capacity for rapid proliferation. However, none of these studies explicitly elucidate the specific function that cytoophidia undertake in cell proliferation. The absence of CTPS redundancy provides *S. pombe* with a distinct advantage as a model organism for investigating the function of cytoophidia. In comparison to humans and *S. cerevisiae*, which possess two CTPS isoforms (CTPS1 and CTPS2 in humans; URA7 and URA8 in *S. cerevisiae*), *S. pombe* only harbors one CTPS isoform encoded by a single locus *cts1* on chromosome I.

In this study, we utilize *S. pombe* as a model system to investigate the role of CTPS cytoophidia in cellular proliferation. Our results demonstrate that the absence of cytoophidia leads to a delay in the G2 phase of the cell cycle and an increase in cell size. Interestingly, there exists a mutual dependence between cytoophidia and CTPS protein levels; specifically, loss of cytoophidia decreases protein levels while protein levels influence cytoophidia formation. The expression levels of multiple genes associated with cell growth and proliferation are reduced in strains lacking cytoophidia. Overexpression of SLM9, a histone transcription regulator involved in regulating CDC2 via WEE1, mitigates the prolongation of the cell cycle and enlargement of cell size caused by loss-filament mutant of CTPS. These findings provide valuable insights into the regulatory roles played by cytoophidia during cellular proliferation.

## 2. Results

### 2.1. Cytoophidia are observed exclusively during the log phase of *S. pombe*

The growth of *S. pombe* cells can be divided into four distinct stages: lag phase, exponential phase, stationary phase, and death phase, and each characterized by unique metabolic conditions. We previously reported that cytoophidia regulated by the metabolic conditions within *S. pombe* [26]. Utilizing the CTPS-YFP tagged strain (CTPS-YFP) previously established in our laboratory, we observed a high abundance of cytoophidia during the logarithmic phase, present in over 90% of fission yeast cells (Fig.1A and C), which subsequently disappeared during the stationary phase (Fig. 1B and C). Furthermore, compared with the exponential phase, there was a decrease in CTPS protein levels during the stationary phase while CTPS RNA levels remained unchanged (Fig.1D, E, and F). These findings indicate that cytoophidia are exclusively present during the logarithmic growth stage of fission yeast and suggest their potential importance in cell proliferation.

**Figure 1.**
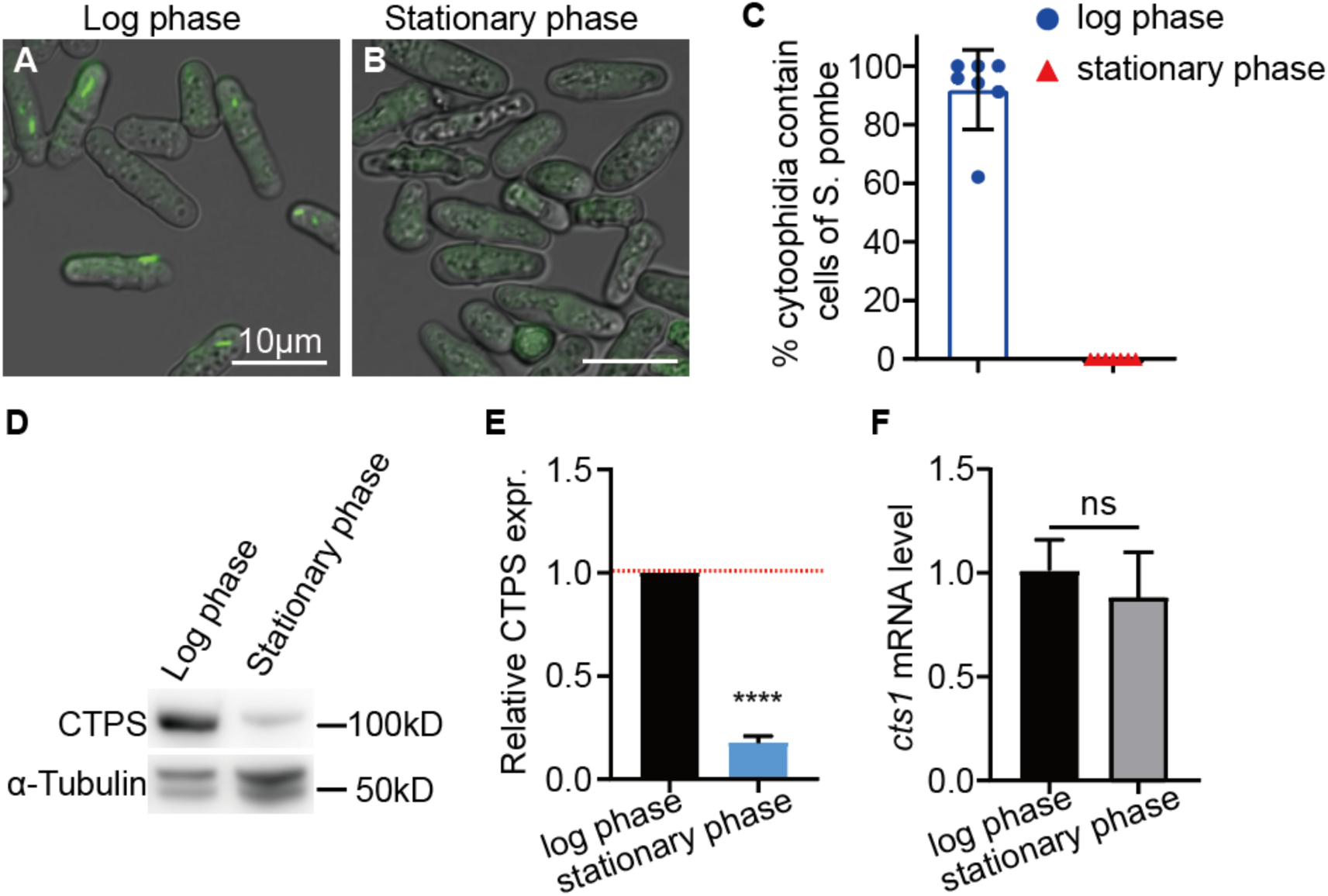
Cytoophidia present only in the log phase of *S. pombe.* **(A-B)** The CTPS protein form the cytoophidia at log phase in fission yeast (A). However, in stationary phase, cytoophidia are disappeared (B). Scale bars, 10 μm. **(C)** Statistics on the percentage of cells containing cytoophidia in fission yeast at log phase and stationary phase. **(D-E)** Western blotting analysis of CTPS proteins from CTPS-YFP tagged fission yeast at log and stationary phase. Anti-YFP antibody was used. Alpha-tubulin was used as an internal control (D). Quantification of the CTPS protein relative expression levels in D (E). (3 biological replicates). **(F)** Levels of CTPS mRNA expression as detected by qPCR in fission yeast at log and stationary phase. Values were normalized against 18s rRNA expression. (3 biological replicates). All values are the mean ± S.E.M. ****p < 0.0001, ns, no significant difference by Student’s t test.

### 2.2. Loss-filament mutant of CTPS impacts the cell cycle and size of *S. pombe*

To investigate the role of cytoophidia in fission yeast, we specifically disrupted the polymerization of CTPS to form cytoophidia without affecting their enzymatic functions. Previous studies have demonstrated that the histidine amino acid at position 355 (His^355^) in human and *Drosophila*, as well as His^360^ in *S. cerevisiae*, is crucial for CTPS polymerization [17,21]. By aligning the amino-acid sequences of both *S. cerevisiae* and *S. pombe* CTPS proteins, we observed a similarity between His^359^ in *S. pombe* CTPS and His^360^ in *S. cerevisiae* (Fig. 2A). Therefore, we hypothesized that the replacement of His^359^ with Ala^359^, an uncharged and smaller amino acid compared with histidine, would disrupt the polymerization ability of the *S. pombe* CTPS protein like that observed in humans, *Drosophila*, and *S. cerevisiae*.

**Figure 2.**
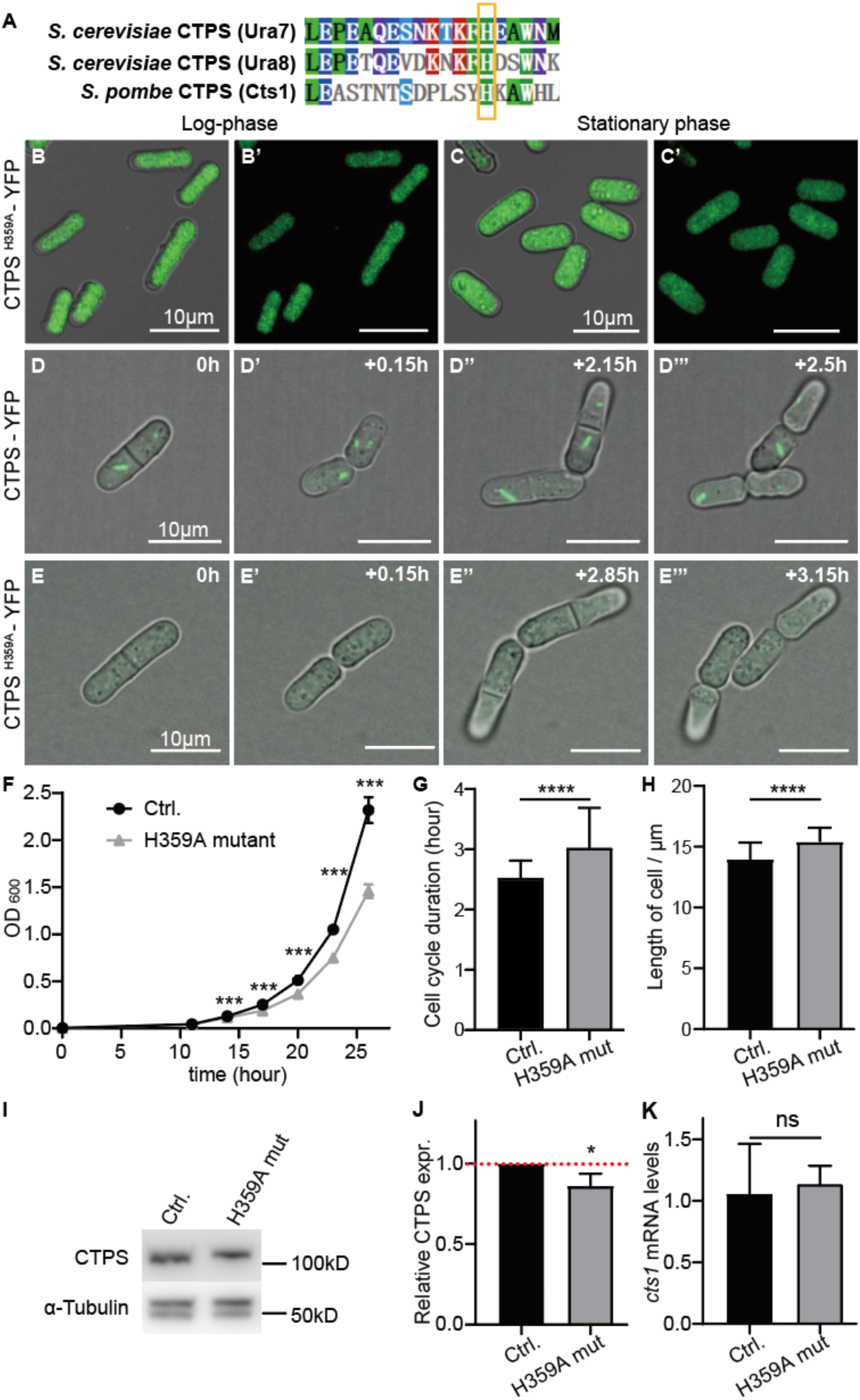
Loss-filament mutant of CTPS affects the cell cycle and size in *S. pombe*. **(A)** Protein sequence alignment between ura7, ura8, and cts1, and found the histidine at site 359 of cts1 is similar to that at site 360 of ura7, which has been reported to mutate histidine to alanine without affecting enzyme activity but preventing cytoophidia formation in S. *cerevisiae* [21]. **(B-C’)** Photos of the histidine at site 359 of cts1 mutated to alanine (H359A mut) in *S. pombe.* Cytoophidia were not formed at both log phase (B, B’) and stationary phase (C, C’). Scale bars, 10 μm. **(D-E’’’)** Live cell imaging photos of CTPS-YFP and CTPS^H359A^-YFP strains at the log phase, show that cytoophidia were formed at the log phase in CTPS-YFP strain (G-G’’’), and not formed in CTPS^H359A^-YFP strain (H-H’’’). Green: endogenous CTPS. Scale bars, 10 μm. **(F)** Growth curve of CTPS-YFP, as control and CTPS^H359A^-YFP strains. CTPS^H359A^-YFP strain grows slower than control. (3 biological replicates). **(G-H)** The cell cycle duration and cell length obtained from the data of live cell imaging showed that CTPS^H359A^-YFP strain had a longer cell cycle (G) and cell length (H) than the control strain (>86 cells were manually measured per strain). (3 biological replicates). **(I-J)** Western blotting analysis of CTPS proteins from CTPS-YFP and CTPS^H359A^-YFP strains at log and stationary phase. Anti-YFP antibody was used. Alpha-tubulin was used as an internal control (I). Quantification of the CTPS protein relative expression levels in I (J). (3 biological replicates). **(K)** Levels of CTPS mRNA expression as detected by qPCR at log phase. Values were normalized against 18s rRNA expression. (3 biological replicates). All values are the mean ± S.E.M. ****P< 0.0001, ***P< 0.001, *P< 0.05, ns, no significant difference by Student’s t test.

Consequently, we generated a point mutant where His^359^ was replaced by Ala^359^ (refered to as H359A) within the endogenous CTPS protein of *S. pombe.* Additionally, we fused YFP at the C-terminus of the mutant CTPS for easy detection (CTPS^H359A^-YFP). By extracting genomic DNA from the H359A mutant fission yeast, we confirmed successful introduction of the intended point mutation (Supplemental Fig. 1). The CTPS H359A mutant fission yeast was viable, but did not form cytoophidia (Fig. 2B-C’).

Furthermore, measurement via the OD_600_ value showed that the CTPS H359A mutant strain exhibited a slower growth rate compared with the CTPS-YFP tagged strain, which served as the control group (Fig. 2F). Live cell imaging data showed that the CTPS H359A mutant strain had an extended cell cycle and increased cell length in comparison to the control (Fig. 2D-E’’’, G and H). Interestingly, western blotting analysis indicated reduced levels of CTPS protein in the H359A mutant strain when compared with the control strain (Fig. 2I and J), while real-time PCR showed similar levels of CTPS RNA between them (Fig. 2K).

Combined with the data of our previous studies [17,19], these findings suggest that cytoophidia formation is crucial for maintaining adequate levels of CTPS protein in *S. pombe*, potentially through polymerization to ensure protein stability; they also indicate that failure to form CTPS cytoophidia leads to elongated cell cycles and influences cell size.

### 2.3. Both CTPS protein levels and cytoophidia affect the cell cycle and size of *S. pombe*

To understand whether the cytoophidia and the CTPS protein levels influenced the cell cycle and size. First, we disrupted filament formation without reducing its protein expression levels. We created a CTPS ^H359A-OE^ strain that overexpressed a mutant form of CTPS (H359A) tagged with mCherry at its C-terminus. we also generated a CTPS^WT-OE^ strain that overexpressed wild-type CTPS tagged with mCherry, as well as a strain expressing only mCherry as the wild-type control. The strains were confirmed by real-time PCR and western blotting (Fig. 3D and E).

**Figure 3.**
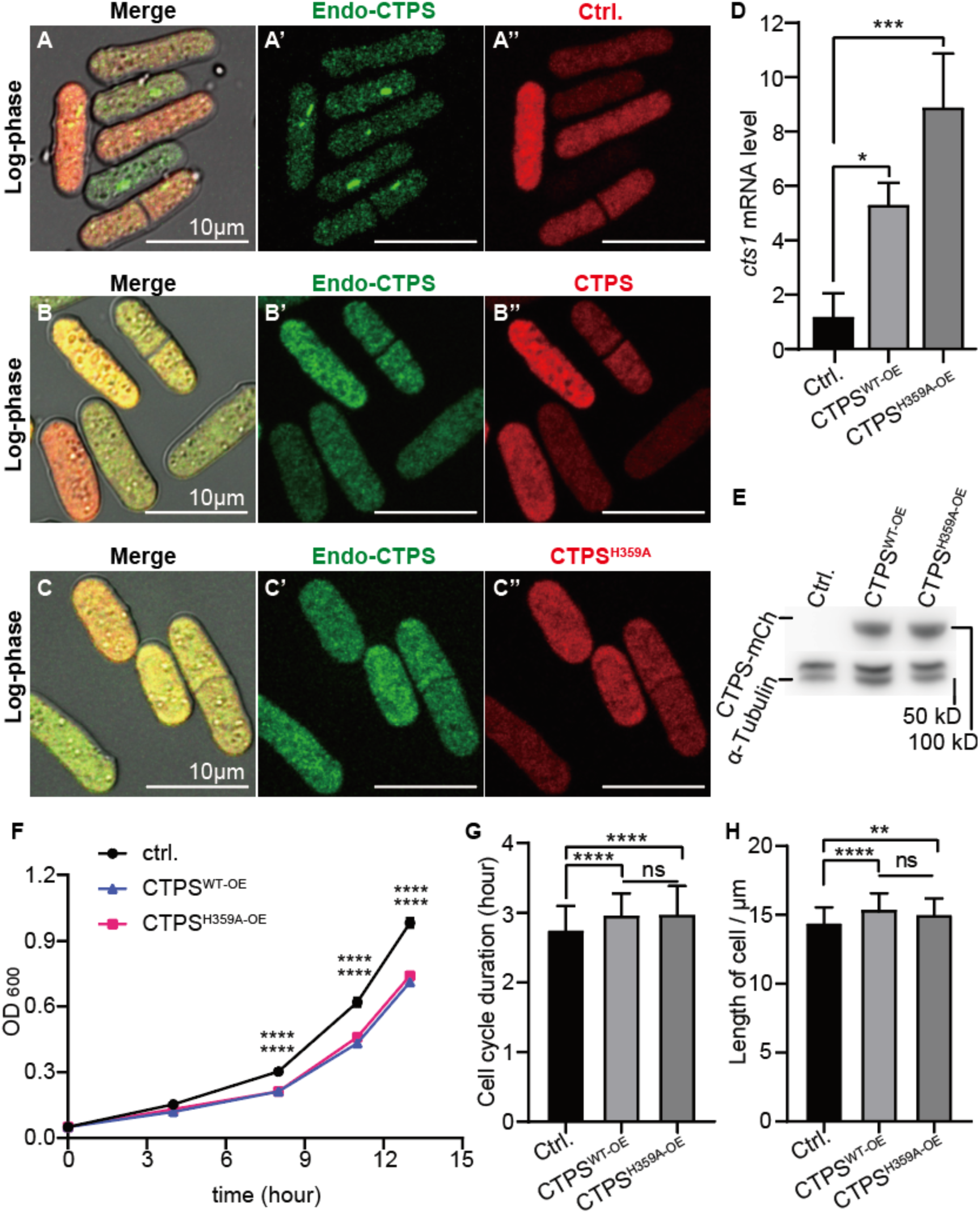
CTPS protein levels influence cytoophidia formation, cell cycle and size in *S. pombe*. **(A-C’’)** Photograph of CTPS-YFP strains that overexpression mCherry (A-A’’), CTPS-mCherry (CTPS^WT-OE^, B-B’’), and CTPS^H359A^-mCherry (CTPS^H359A-OE^, C-C’’) at the log phase. Cytoophidia did not appear in overexpression CTPS and CTPS^H359A^ strains compared with wild-type control group. Scale bars, 10 μm. **(D)** Levels of CTPS mRNA expression as detected by qPCR in control, OE CTPS, and OE CTPS^H359A^ strains at log phase. Values were normalized against 18s rRNA expression. One-way ANOVA was used for statistical analysis. **(E)** Western blotting analysis of CTPS proteins from control, OE CTPS, and OE CTPS^H359A^strains at log phase. Anti-mCherry antibody was used. Alpha-tubulin was used as an internal control. **(F)** Growth curve of control, OE CTPS, and OE CTPS^H359A^strains. OE CTPS and OE CTPS^H359A^ strains grow slower than control. Unpaired Student’s t test was used for statistical analysis. (3 biological replicates). **(G-H)** The cell cycle duration and cell length obtained from the data of live cell imaging showed that OE CTPS and OE CTPS^H359A^ strains had a longer cell cycle (G) and cell length (H) than control. (>85 cells were manually measured per strain). One-way ANOVA was used for statistical analysis. (2 biological replicates). All values are the mean ± S.E.M. ****P< 0.0001, ***P< 0.001, **P< 0.01, *P< 0.05, ns, no significant difference.

In both the log and stationary phases, the presence of cytoophidia was not observed in either the CTPS ^H359A-OE^ or the CTPS ^H359A^ strain, suggesting that the H359A mutation can act dominantly to inhibit cytoophidia formation (Fig. 3C-C’’ and Supplemental Fig. 3C-C’’). Surprisingly, even in the presence of overexpressed wild-type CTPS in the CTPS^WT-OE^ strain, cytoophidia did not appear during both log phase (Fig. 3A-B’’) and stationary phase (Supplemental Fig. 2A-B’’). Additionally, based on OD_600_ measurements from growth curves, both the CTPS^WT-OE^ and CTS^H359A-OE^ strains exhibited slower growth rates compared with the wild-type control (Fig. 3F).

Live cell imaging data revealed that both these strains had longer cell cycles and increased cell length compared with the wild-type control (Fig. 3G-H). These observations were consistent with those seen in the CTS^H359A^ strain (Fig. 2I-G-H), indicating an influence of both cytoophidia formation and CTPS protein levels on cell proliferation rate.

Second, to investigate whether the decreasing CTPS protein levels influenced the formation of cytoophidia and cell proliferation. We knock down the CTPS using the CRISPRi method based on the CRISPR-dCas9 system [27]. The expression of dCas9 protein was regulated by the nmt1-41 promoter, which induced transcription upon thiamine removal from the medium (Supplemental Fig. 3A). Five sgRNAs were designed to target the *cts1* gene encoding CTPS protein in fission yeast (Supplemental Fig. 3B). Each of these sgRNAs was transformed into a CTPS-YFP tagged fission yeast strain, while a non-homologous served as a control group. Western blotting analysis confirmed the expression of dCas9 protein in all strains (Fig. 4A and C), and showed that CTPS protein levels were reduced in strains containing sgRNA2, sgRNA4, and sgRNA5 compared with the control group after thiamine removal (Fig. 4A and B). However, no significant changes in CTPS protein levels were observed in cells containing sgRNA1 and sgRNA3 (Fig. 4A and B).

**Figure 4.**
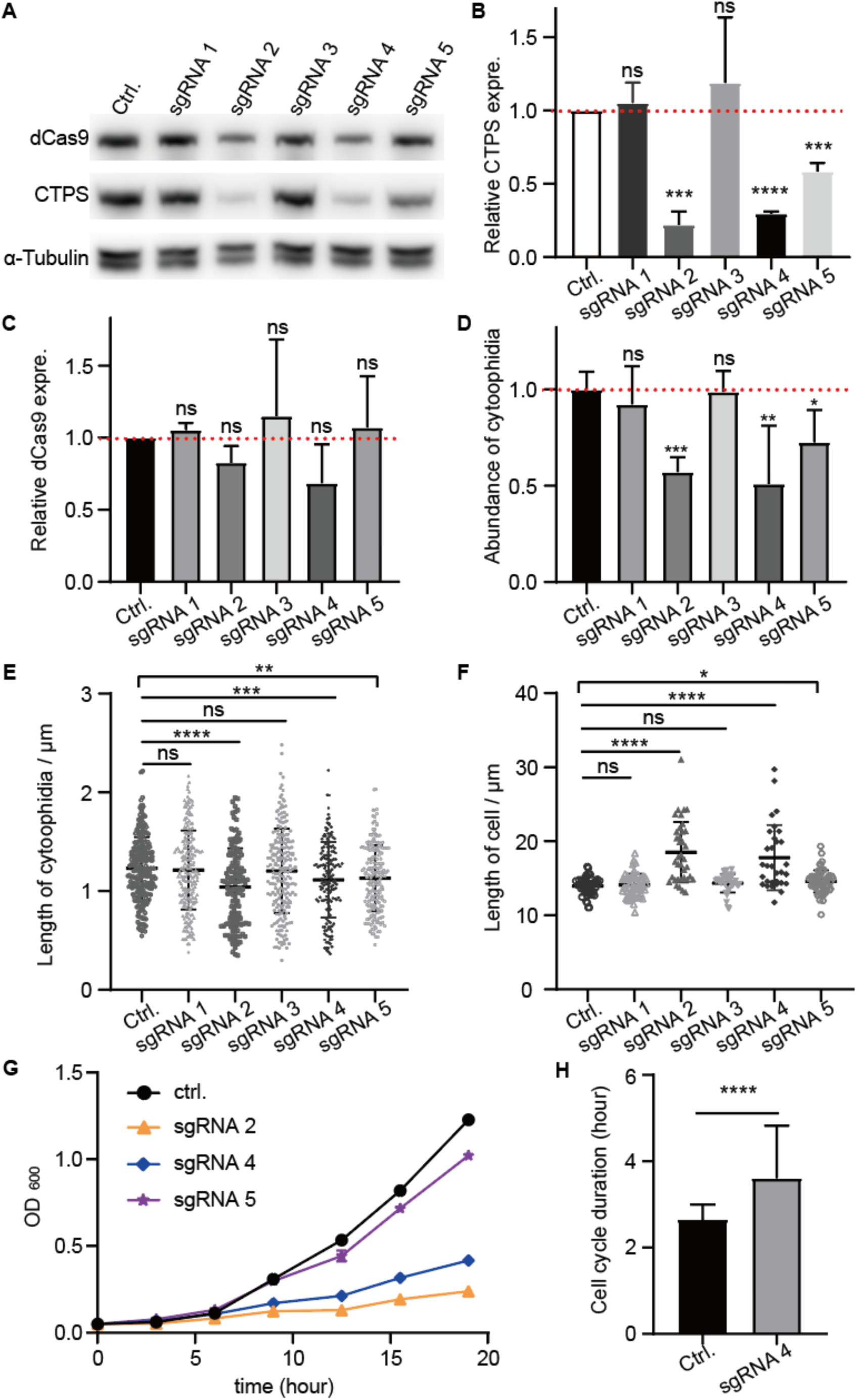
CTPS reduction affects cytoophidia formation, elongates cell cycle and increases cell size. **(A)** Immunoblotting analysis of CTPS and dCas9 proteins from the CTPS-YFP strains which expressed the pdCas9 plasmid carried nonsense sgRNA (as control group), sgRNA1, sgRNAs2, sgRAN3, sgRAN4, sgRAN5, respectively, at log phase. Anti-YFP and anti-flag antibodies were used. Alpha-tubulin was used as an internal control. **(B-C)** Quantification of the CTPS protein (B) and dCas9 protein (C) relative expression levels in A (3 biological replicates). **(D-F)** Quantification of the numbers of cytoophidia (D), length of cytoophidia (E), and cell length (F) in the strains which contained sgRNA and dCas9. The relative value was normalized with control. (>265 cells were manually measured, 2 biological replicates) **(G)** The growth curve of the strains contained nonsense sgRNA (as control group), sgRNA2, sgRNAs4, and sgRAN5. Showed that knockdown CTPS strains grow slower than control. (3 biological replicates). **(H)** The cell cycle duration obtained from the data of live cell imaging showed that knockdown CTPS strains had a longer cell cycle than control. (>230 cells were manually measured, 2 biological replicates). All values are the mean ± S.E.M. ****P< 0.0001, ***P< 0.001, **P< 0.01, *P< 0.05, ns, no significant difference by Student’s t test.

Knocking down CTPS resulted in decreased proportions of cytoophidia-forming cells with shorter cytoophidia lengths observed in cells targeted by sgRNA2, sgRNA4, and sgRNA5 compared with the control group; whereas no changes were seen for cells containing sgRNA1 or sgRNA3 (Fig. 4D and E, Supplemental Fig. 3C-H). Measurement via OD_600_ values revealed slower cell growth rates for CTPS knockdown strains (sgRNA2, sgRNA4, and sgrNA5) compared with the control group (Fig. 4G). The growth rate of sgRNA targeted strains was sgRNA5>sgRNA4>sgRNA2, which the CTPS protein knockdown levels increased sequentially (Fig. 4G), and the cell length of these strains was also longer than the control (Fig. 4F). Furthermore, growth curve data indicated an extended cell cycle duration for CTPS knockdown cells relative to controls (Fig. 4H).

The findings demonstrate that the modulation of CTPS protein levels has an impact on cytoophidia formation and cellular proliferation. Moreover, both CTPS protein levels and cytoophidia in *S. pombe* exert an influence on cell cycle progression and size regulation.

### 2.4. Loss-filaments mutant of CTPS extends the duration of the G2 phase in *S. pombe*

The CTPS^H359A^-YFP strain exhibited a longer cell cycle duration compared with the CTPS-YFP strain. Therefore, we aimed to investigate which specific phase of the cell cycle (S, M, G1, or G2) was affected by this mutation. In fission yeast, the nuclear cell cycle is divided into distinct phases: G1, S, G2, and M. Following mitosis, newly replicated nuclei enter the subsequent cell cycle and undergo G1 and S phases before completing cytokinesis from the previous cycle [28,29]. This differs from mitosis in mammals and budding yeast.

First, we used hydroxyurea (HU) to induce cell cycle synchronization at the beginning of the S phase, the fission yeast was treated with HU for 4 hours followed by a release period of 3 hours. Based on literature summarizing mitosis in fission yeast (Fig. 5D), we performed microscopic imaging of HU-treated cells and stained their nuclei with DAPI dye. Analyzing the state and morphology of the nucleus in these images and statistically evaluating data (Fig. 5A), we observed an increased proportion of cells in the G2 phase within the CTPS^H359A^-YFP strain compared with the CTPS-YFP strain (Fig. 5B). The findings suggest that there is a prolonged duration of G2 phase during the cell cycle progression in the CTPS^H359A^-YFP strain.

**Figure 5.**
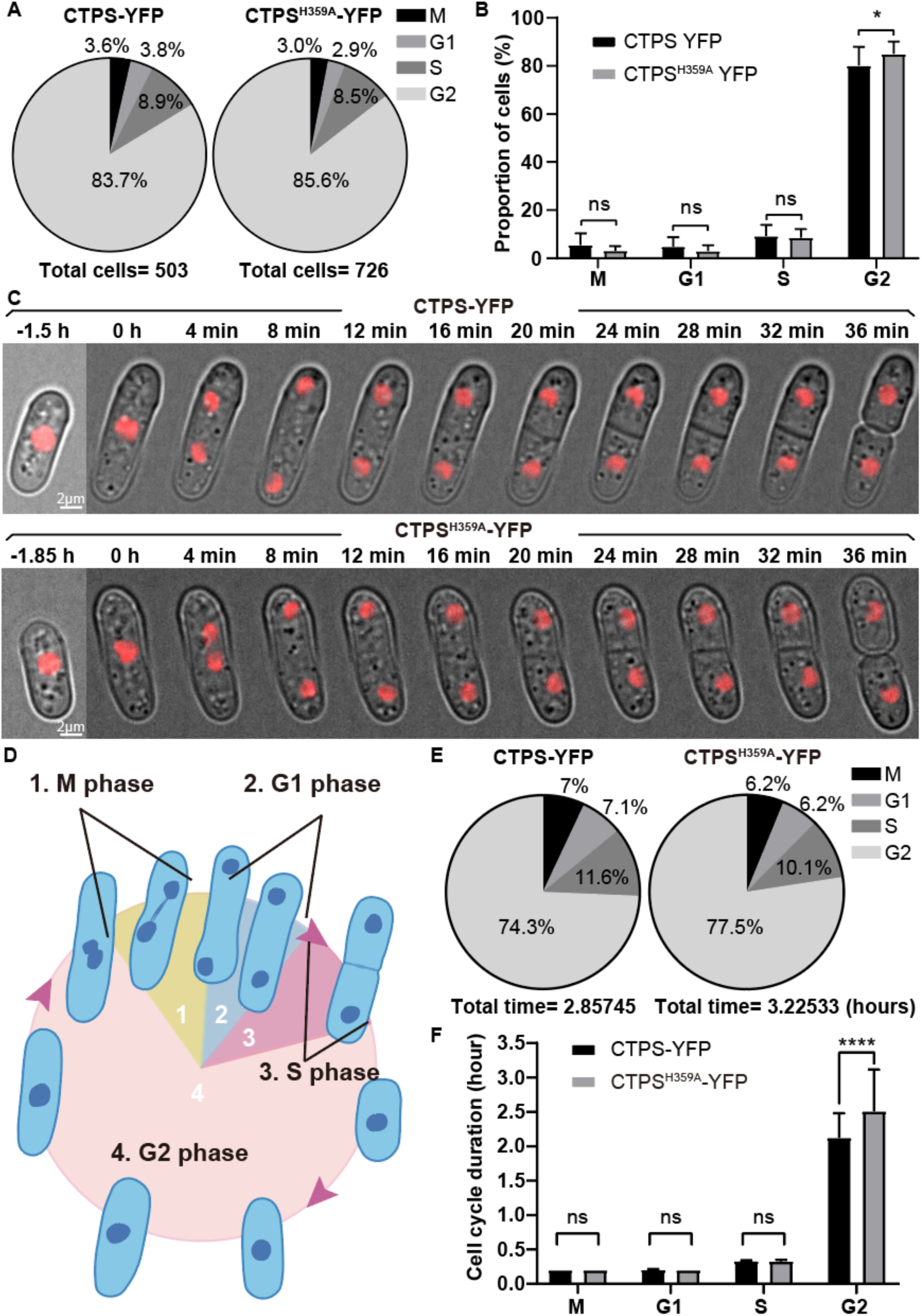
Loss-filament mutant of CTPS prolongs the G2 phase in *S. pombe*. **(A)** The proportion of cells in M, G1, S, and G2 phase of the whole cell cycle, from the fission yeast that was treated with hydroxyurea (HU). **(B)** Quantification of the proportion of cells in M, G1, S, and G2 phase in **A**. CTPS^H359A^-YFP strain had a higher proportion of cells than CTPS-YFP strain at G2 phase. **(C)** Live cell imaging photos of H2B-mCherry expressed fission yeast, showed that CTPS^H359A^-YFP strain grows slower than controls during period of cell volume growth which is in G2 phase. Scale bars, 2 μm. **(D)** Partitioning diagram of M, G1, S, and G2 phase during fission yeast mitosis. **(E)** The time proportion of M, G1, S, and G2 in the whole cell cycle from the data of live cell imaging, >100 cells were manually measured per strain. **(F)** Quantification of the time proportion of M, G1, S, and G2 in the whole cell cycle in **E**. Data showed that CTPS^H359A^-YFP strain had a longer G2 phase than CTPS-YFP strain. All values are the means ± S.E.M. ****p < 0.0001, *p < 0.05, ns, no significant difference by Student’s t test.

To validate the aforementioned inference, we conducted live cell imaging to observe cell division in fission yeast. To label the nucleus, H2B-mCherry was expressed in both the CTPS^H359A^-YFP strain and CTPS-YFP strain. Our results showed that H2B-mCherry expression was observed in both strains, with CTPS-YFP serving as a control group (Fig. 5C).

Following an analysis of the live cell imaging data and observation of nuclear state and morphology, we statistically determined the occurrence time proportions of G1, S, G2, and M phases throughout the entire cell cycle (Fig. 5E), based on previously summarized literature regarding mitosis in fission yeast. Compared with the control group, we found that the G2 phase was significantly prolonged in cells carrying the H359A mutation while no significant differences were observed for G1, S, and M phases (Fig. 5F). These findings provide evidence that loss-filament mutant of CTPS result in an extended G2 phase.

### 2.5. Loss-filament mutant of CTPS perturbs G2/M transition and the expression of cell growth related genes

The CTPS^H359A^-YFP strain exhibits larger cell sizes and a longer G2 phase compared with the CTPS-YFP strain, aiming to investigate whether the loss of filaments mutant of CTPS affects the expression of genes related to cell growth and G2/M transition. We assessed the expression levels of relevant genes using real-time PCR in fission yeast at the logarithmic phase, including G2/M transition regulators (*wee1*, *cdc25*, *cdr1*, *cdr2*, *cdc13*, *fin1*, *blt1*, and *slm9*) and cell growth regulators (*tea1*, *pom1*, *sty1*, *pop3*, *tor1*, and *tor2*). Our data revealed that in the CTPS^H359A^-YFP strain, the gene expression levels of *cdc25*, *cdr1*, *cdc13*, *fin1*, *blt1*, *slm9*, *pop3*, *tor1*, and *tor2* were decreased compared with those in the CTPS-YFP strain (Fig.6). This suggests that loss-filaments mutant of CTPS influences the expression of genes associated with cell growth and G2/M transition.

**Figure 6.**
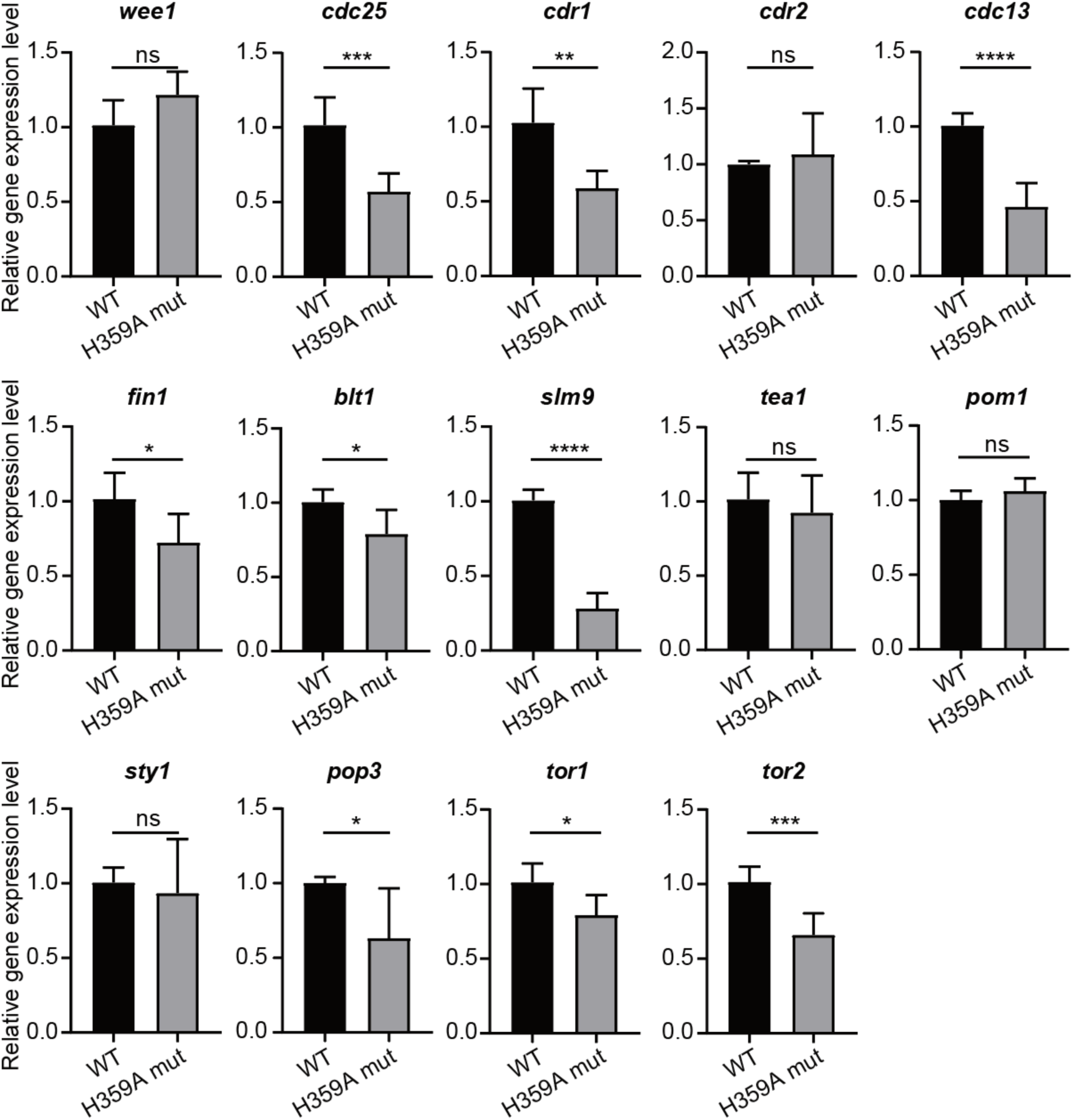
Loss-filament mutant of CTPS perturbs G2/M transition and the expression of cell growth-related genes. RNA levels of some regulators of cell size and cell cycle in G2/M transition in CTPS^H359A^-YFP and CTPS-YFP strains at the log phase detected by real-time PCR. Including G2/M transition regulators: *wee1*, *cdc25*, *cdr1*, *cdr2*, *cdc13*, *fin1*, *blt1*, and *slm9*; cell growth regulators: *tea1*, *pom1*, *sty1*, *pop3*, *tor1*, and *tor2*. All values are the means ± S.E.M. ****p < 0.0001, ***p < 0.001, **p < 0.01, *P< 0.05, ns, no significant difference by Student’s t test. (3 biological replicates)

### 2.6. Overexpressing *slm9* alleviates phenotypes caused by loss-filament mutant of CTPS

The histone transcription regulator SLM9 has been previously shown to regulate CDC2 activity through WEE1, and *slm9* deficiency strain exhibit a G2 cell cycle delay [30]. Immunoaffinity purifications of SLM9 have revealed the co-purification of CTS1 with SLM9 proteins [31]. Additionally, our previous study reported that *slm9* knockout affects cytoophidia, increases cell volume, and slows down cell growth rate in *S. pombe* [32].

In this study, we found that the loss-filament mutant of CTPS reduces the expression level of *slm9*. These findings suggest that SLM9 is involved in G2 prolongation caused by the loss-filament mutant of CTPS. Therefore, we investigated whether overexpressing *slm9* could rescue prolonged cell cycle and enlarged cell size in the CTPS^H359A^-YFP strain. We overexpressed *slm9*-mCherry in the CTPS^H359A^-YFP strain, the SLM9 is localized to the nucleus, referred to as OE-*slm9* strain (Fig. 7C-C’’, Supplemental Fig. 4A-A’’). Control strains were also generated: control.1 and control.2 strains only overexpressed mCherry in the CTPS-YFP and CTPS^H359A^-YFP strains respectively (Fig. 7A-B’’, Supplemental Fig. 4B-C’’). The *slm9* expression levels in OE-*slm9* strain were confirmed by real-time PCR compared with the control.1 strain (Fig. 7D).

**Figure 7.**
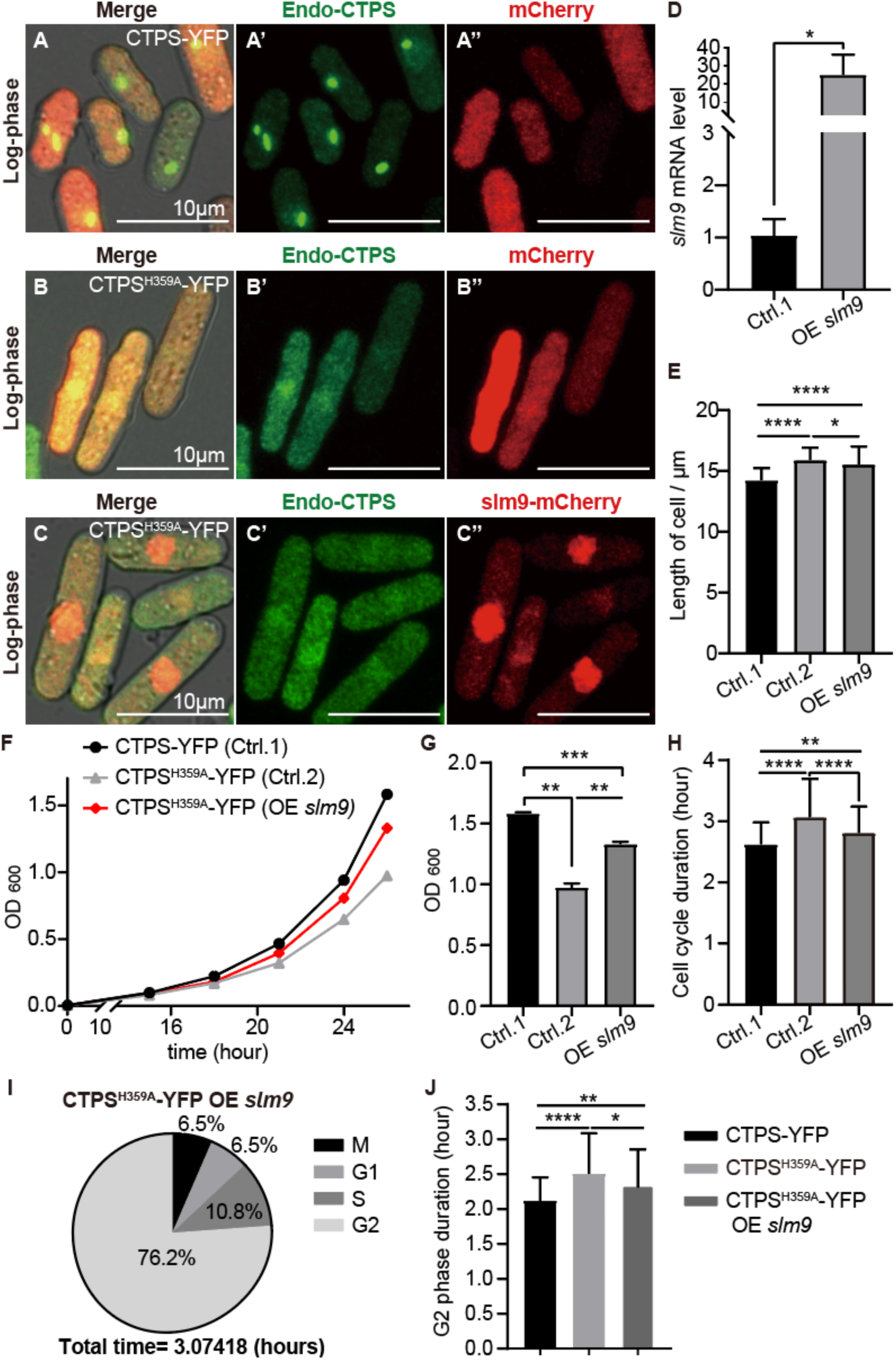
Overexpressing *slm9* alleviates cell cycle prolongation caused by loss-filament mutant of CTPS. **(A-C’’)** Photograph of CTPS-YFP strain that overexpressed only mCherry (named Control.1, A-A’’), CTPS^H359A^-YFP strain that overexpressed only mCherry (named Control.2, B-B’’), and CTPS^H359A^-YFP strain that overexpressed slm9-mCherry, SLM9 localized to the nucleus (named OE-*slm9*, C-C’’) at the log phase. Scale bars, 10 μm. **(D)** Levels of *slm9* mRNA expression as detected by qPCR in control.1 and OE-*slm9* strains at log phase. Values were normalized against 18s rRNA expression. Unpaired Student’s t test was used for statistical analysis. **(E)** Cell length obtained from the data of live cell imaging showed that OE *slm9* in CTPS^H359A^-YFP strain had a shorter cell length than control.2 strain, but longer than control.1 strain. (>120 cells were manually measured per strain). One-way ANOVA was used for statistical analysis. **(F)** Growth curve of control.1, control.2, and OE-*slm9* strains. OE *slm9* strain grows faster than control.2 strain, but slower than control.1 strain. (3 biological replicates). **(G)** The OD_600_ value of control.1, control.2, and OE-*slm9* strains at 26h in (F), statistical analysis by one-way ANOVA, showed that OE-*slm9* strain grows faster than loss-filament mutant strain, but slower than normal unmutated strain. **(H)** The cell cycle duration obtained from the data of live cell imaging showed that the cell cycle of the OE-*slm9* strain was shorter than control.2 but longer than control.1. (>120 cells were manually measured per strain). One-way ANOVA was used for statistical analysis. **(I)** The time proportion of M, G1, S, and G2 in the whole cell cycle from the live cell imaging data of OE *slm9* strain (>100 cells were manually measured per strain). **(J)** Compared the G2 phase duration of the CTPS-YFP, CTPS^H359A^-YFP, and OE-slm9 strains, showed that G2 phase of OE-*slm9* strain was shorter than CTPS^H359A^-YFP strain, but longer than CTPS-YFP strain. Statistical analysis by one-way ANOVA. (>100 cells were manually measured per strain). All values are the mean ± S.E.M. ****P< 0.0001, ***P< 0.001, **P< 0.01, *P< 0.05.

Growth curves measured via OD600 values showed that the OE-slm9 strain grew faster than the control.2 but slower than control .1 (Fig. 7F). A statistically significant difference was observed at 26h when comparing OD600 values between these strains; indicating that OE-*slm9* grows faster than loss-filament mutants but slower than normal unmutated strains (Fig. 7G). Furthermore, analysis of live cell imaging data revealed that the cell cycle duration for OE-slm9 was shorter than control .2 but longer than control. 1 (Fig. 7H). Cell length measurement showed that OE-*slm9* strain exhibited a shorter cell length compared with the control.2 strain, but longer than the control.1 strain (Fig. 7E). Additionally, a statistical analysis was performed on the proportion of occurrence time for G1, S, G2, and M phases throughout the entire cell cycle (Fig. 7I).

Comparisons of the duration of the G2 phase among CTPS-YFP, CTPS^H359A^-YFP, and OE-slm9 strains revealed that in the OE-slm9 strain, the G2 phase was shorter compared with the CTPS^H359A^-YFP strain but longer than in the CTPS-YFP strain. However, no significant differences were observed regarding the durations of G1, S, and M phases (data not shown). These results indicated that overexpression of *slm9* alleviates cell cycle prolongation at the G2 phase and cell size enlargement caused by loss-filament mutant of CTPS.

## 3. Discussion

The presence of cytoophidia in a wide range of species, from prokaryotes to eukaryotes, suggests that it likely serves an important biological function. Although it has been observed in highly proliferative cells such as human hepatocellular carcinoma and mouse thymus T cells [5,10], indicating its involvement in cell proliferation, the exact role of cytoophidia remains enigmatic. Therefore, we aimed to investigate the impact of cytoophidia on cell proliferation and elucidate its underlying mechanisms.

In this study, using *S. pombe* as a model organism, we demonstrate that loss-filament mutant of CTPS causes a delay in the G2 phase of the cell cycle and results in enlarged cell size during *S. pombe* proliferation. These findings are consistent with previous reports in budding yeast where dysregulated assembly of CTPS leads to impaired growth [21]. Furthermore, expression levels of various genes associated with G2/M transition during cell proliferation were found to be decreased in the CTPS^H359A^ strain. Collectively, these results indicate that cytoophidia play a role in regulating cell proliferation by participating in G2 phase regulation.

Studies have demonstrated that substitution of the wild-type CTPS protein with a mutant form, in which H^355^ is converted to A^355^ in humans and *Drosophila*, and H^360^ is converted to A^360^ in budding yeast, leads to a failure in cytoophidia formation without affecting the tetramer structure of CTPS, which is essential for its activity [14,33]. By aligning the amino acid sequences of CTPS proteins from both *S. cerevisiae* and *S. pombe*, we observed that conversion of amino acid H^359^ to A^359^ failed cytoophidia formation in fission yeast as expected.

Previous studies have shown that cytoophidia stabilize the CTPS protein by prolonging its half-life; specifically, the H355A mutant reduced CTPS protein levels in human cancer cells and *Drosophila* adipocytes [17,19]. In our study, we observed a slight decrease in CTPS protein levels in the CTPS^H359A^-YFP strain, similar to what was observed with the H355A mutant in human cells and *Drosophila* adipocytes. Interestingly, we found that loss of cytoophidia decreased CTPS protein levels; furthermore, both knockdown and overexpression of CTPS protein influenced cytoophidia formation. These findings suggest mutual regulation between cytoophidia and CTPS protein levels within cells.

Our previous study showed that histone transcription regulator SLM9 is required for cytoophidia biogenesis, Knockout of *slm9* resulted in a decrease in the percentage of cells containing cytoophidia and mitotic cells, while increasing the length of both cytoophidia and cells in *S. pombe* [32]. In our study, loss-filament mutants of CTPS exhibited reduced expression levels of *slm9*, leading to elongation of cell length and delayed cell cycle progression. The observed phenotypes were similar to those observed in *slm9* knockout strains, suggesting a mutual regulation between *slm9* and cytoophidia.

Furthermore, studies have shown that SLM9 regulates CDC2 activity through WEE1, deficiency *slm9* leads to G2 cell cycle delay; immunoaffinity purifications of SLM9 have shown co-purification with CTS1 proteins [30,31]. Our study showed that overexpression of *slm9* partially rescue the elongation of G2 phase and cell enlargement induced by loss-filament mutants of CTPS. These findings suggest the involvement of slm9 in the regulation of cytoophidia on cell proliferation.

Since the discovery of cytoophidia in 2010, extensive research has been conducted to elucidate the biological significance of their existence. In this study, we have identified that cytoophidia exert an influence on cell proliferation by modulating the G2 phase, with slm9 also implicated and a potential regulatory relationship between slm9 and cytoophidia (Fig. 8). However, the precise mechanism through which cytoophidia regulate cell proliferation remains elusive. Our investigation may provide valuable insights into unraveling this mechanism. This will be a direction of exploration, moreover, we await future studies aimed at the investigation of more biological functions of the cytoophidia in different species.

**Figure 8.**
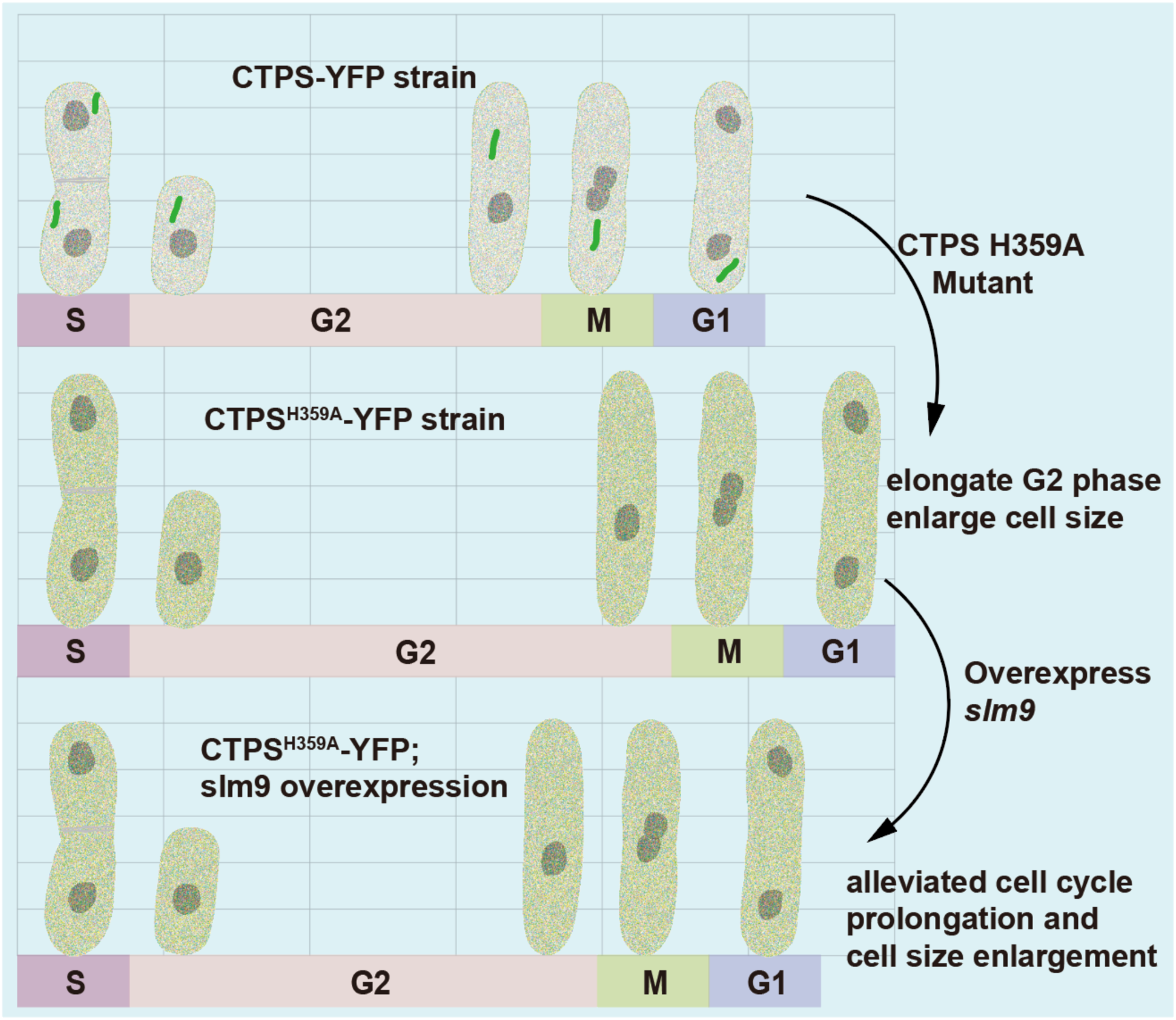
Schematic model for the effect of cytoophidia on cell growth and size. Loss of cytoophidia leads to a delay G2 phase in cell cycle and enlarges cell size. Overexpressing slm9 alleviates cell cycle prolongation and cell size enlargement caused by loss-filament mutant of CTPS in *S. pombe*.

## 4. Materials and methods

### 4.1. Yeast strain and culture medium

Fission yeast cts1-YFP strain was constructed as previously described [7], specifically, the wild type *S. pombe* strain: h-; ade6-M216, leu1-32, ura4-D18 was transformed linearized (SpeI) pSMUY2-*Ura4*-*cts1*-YFP plasmid. Similarly, the Cts1^H359A^-YFP strain was constructed using a PCR-based approach for gene tagging in normal chromosomal locations in fission yeast [34]. Briefly, forward and reverse primers were designed to be complementary to a 20-bp region on each side of the knock-in cassette. Additionally, flanking sequences of 60 bp that were complementary to the immediate 5’-upstream and 3’-downstream endogenous regions of the cts1 gene were included in each primer (total length of 80-90 bp for each primer). These primers were then used in a PCR reaction with the *cts1*-H359A-YFP containing plasmid as template to amplify the cassette. The resulting PCR product was purified and approximately 1-2 µg was transformed into fission yeast using the lithium acetate method [34]. The primers used in this study are listed in **Table S2**. Strains carrying plasmids with overexpressed genes were also prepared using the lithium acetate method. A comprehensive list of all viable *S. pombe* strains can be found in **Table S1**. Fission yeast was cultured in standard rich media (YE4S; complete medium enriched with supplements including leucine, uracil, histidine, and adenine at a concentration of 100 µM) or Edinburgh minimal medium 2 (EMM2) containing 3% glucose and supplemented with necessary amino acids (leucine, uracil, histidine, and adenine at a concentration of 100 µM) for yeast survival at either 30 ℃ or 32 ℃. For plate culture, an additional 2% agar was added and incubated at a temperature of 30 degrees for 3-5 days.

### 4.2. Plasmid construction

The overexpression plasmids were all constructed using the ClonExpress Ultra One Step Cloning kit (Vazyme, Cat. C115-01), genes for the overexpressed plasmids were amplified by PCR from the yeast genome and integrated into corresponding vectors with specific nutritional deficiencies. For example, to construct a plasmid for overexpressing cts1 or cts1^H359A^ genes, fission yeast genomic DNA was used as a template for PCR amplification primers with point mutations designed to obtain the cts1 H359A mutant gene. The cts1 and cts1^H359A^ genes were then inserted into leucine selection-labeled overexpression vectors, while slm9 and Htb1 genes were inserted into adenine-labeled overexpression vectors of fission yeast. All constructed plasmids underwent verification of their DNA sequences by Azenta.

### 4.3. Gene suppression

The *cts1* gene was knocked down using dcas9-mediated CRISPRi. The plasmid pAH237, generously provided by Prof. Tomoyasu Sugiyama, harbors the coding sequence of Cas9 along with a cloning site for the sgRNA and a leucine genetic screening marker. Subsequently, modifications were made to this plasmid to enable the expression of catalytically inactive dCas9 [27]. Briefly, dCas9 eliminates DNA endonuclease activity through mutations D10A and H840A of cas9. The resulting plasmid, which expresses sgRNA and dCas9, is named pdCas9. Targeting sequences (sgRNA) were designed using https://crispr.dbcls.jp/, which searched for specific 20-bp targeting sequences adjacent to PAM sequences (5’-NGG-3’) in the *S. pombe* genome [35]. The constructed sgRNA-pdCas9 plasmids were transferred into fission yeast using the lithium acetate method. Transcription of the dCas9 gene is controlled by an inducible promoter, nmt1-41p, whose transcription is inhibited by 15 µM thiamine in the culture medium. Therefore, fission yeast cells carrying pdCas9 plasmid were grown on the EMM2 plates supplemented with 20 µM thiamine and necessary amino acid for yeast survival. To induce expression of dCas9 in fission yeast, thiamine was removed by washing the cells with sterilized water twice. Cells were then cultured in thiamine-free EMM2 medium at 32 ℃ for approximately 20 h with shaking. A list of all sgRNA oligonucleotides and primers used in dcas9-mediated CRISPRi can be found in **Table S2**.

### 4.4. Growth assays

Cells were cultured in YES or EMM2 supplemented with necessary amino acids for at least two passages to maintain high proliferative activity. The growth curve was plotted by measuring OD_600_ after reviving cells to a robust growth state and diluting them to an initial OD_600_ of ∼0.05 or lower concentrations, such as 0.005. Changes in optical density were measured at corresponding time points during culture. For cell cycle synchronization and release, hydroxyurea (HU) was added to exponentially growing fission yeast at a final concentration of 12 mM, incubated for 4 hours, and removed by centrifugation (1000 g for 5 minutes) at room temperature. The collected sample was washed twice and cultured for three hours to release cells from synchronization. All cells were cultured at 32 ℃.

### 4.5. Cell fixation, image acquisition, and live cell imaging

For the experiment requiring fixed samples, fission yeast cells were collected during either the exponential stage or starvation stage and fixed in 4% paraformaldehyde for 10 minutes at a temperature of either 30 ℃ or 32 ℃ with shaking at 250 rpm. The fixed cells were then washed with PBS and mixed with a ratio of 2:1 with 1.5% agarose before being dropped onto a glass slide. After covering the sample with a coverslip, the film was sealed using nail polish. Images of the fixed samples were acquired using a Zeiss LSM 980 Airyscan2 microscope equipped with a Plan APO 63x/1.40 OIL objective in airyscan mode.

For live cell imaging, fission yeast cells were cultured in glass-bottomed Petri dishes (Thermo scientificTM, Cat. 150460). Specifically, fission yeast was cultured until it reached an exponential state corresponding to an OD_600_ value between 0.5 and 0.8. Then, approximately 0.5µl of culture was placed at the center of each glass-bottomed Petri dish followed by gentle addition of dropwise low-melting agarose solution (1.5%) totaling about 400 µl. After allowing it to stand at room temperature for 10 minutes, the culture medium (about 1 ml) was added. Zeiss Cell Observer SD spinning disk confocal microscope equipped with a 63x OIL objective was used for live cell imaging at 32℃.

### 4.6. Western blotting

Fission yeast protein was extracted using alkaline lysis, followed by western blotting were performed [36]. Briefly, for the protein extraction, 15 ml of exponential stage or starvation stage fission yeast were collected and washed with 1 ml distilled water. The cells were then resuspended in 0.3 ml of distilled water, and treated with an equal volume of 0.6 M NaOH for 10 min at room temperature. After centrifugation and removal of the supernatant, the cells were resuspended in 150 µl of 1x SDS sample buffer (Abclonal, Cat. RM00001) and incubated at 95℃ for 5 min before being briefly centrifuged to collect the supernatant. 15 µl of the supernatant was loaded onto a gradient SDS-polyacrylamide gel (GenScript, Cat. M00654) and electrophoresed at 120 V in regular Tris-glycine buffer. Protein transfer onto polyvinylidene difluoride membrane was performed using a Trans-blot turbo system (Bio-Rad). After blocking with 5% milk, target proteins were detected by using primary antibodies of mouse monoclonal anti-GFP antibody (1: 2000; Roche, Cat. 11814460001), mouse anti-mCherry (1:5000, Abbkine, Cat. Ao2080) and mouse anti-α-Tubulin antibody (1:5000; Sigma, Cat. T6199), followed by secondary antibody Horseradish peroxidase (HRP)-conjugated anti-mouse IgG HRP-linked (1:5000, Cell Signaling, Cat. 7076). Non-saturated bands were quantified on ImageJ (National Institutes of Health) and presented as a ratio relative to the internal reference α-Tubulin. At least two to three biological replicates were quantified.

### 4.7. RNA isolation and RT-PCR

Total RNAs were extracted using TransZol Up Plus RNA Kit (TransGen Biotech, Cat. ER501-01). To disrupt the cell walls of fission yeast, 15 ml log phase cells at the same time point or 5 ml stationary phase cells were collected by centrifugation and resuspended in 1 ml of TransZol Up reagent. Micro glass beads (∼200 µl in volume) were added to the suspension, and the cells were then beaten with the glass beads for 10 min on a Vortex machine (30 s of beating followed by 30 s of resting time). Subsequently, the solution was extracted with 200 µl phenol following the instructions provided in the kit. Equal amounts of RNA templates were reverse transcribed into cDNAs using ABScript III RT Master Mix for qPCR (ABclonal, Cat. RK20429). The resulting cDNAs were utilized in qPCR reactions employing Universal SYBR Green Fast qPCR Mix (ABclonal, Cat. RK21203) on a BIO-RAD CFX Connect Real-Time System according to the manufacturer’s protocol. Relative expression levels were determined using the comparative CT method with normalization against 18s rRNA as an internal control. The primers used for RT-PCR are listed in **Table S3**.

### 4.8. Cell cycle and cell size analysis

ImageJ was utilized for the quantification of cell size, specifically by measuring the cell length of fission yeast. The determination of cell cycle duration was manually performed using ZEN (blue edition) software, which facilitated live cell imaging data analysis and recording of the time required for cell division. Subsequently, statistical analysis was conducted to assess the significance of differences in both cell cycle duration and cell length between various groups, employing One-way ANOVA or Unpaired Student’s t-test.

### 4.9. Quantification and statistical analysis

The sample size for each figure is indicated in the figure legends. Statistical significance between conditions was assessed using Unpaired Student’s t-test. For multiple group comparisons, one-way ANOVA analysis was performed. All statistical analyses were performed in GraphPad Prism, all error bars represent the mean ± standard errors (S.E.M), and significance is denoted as *p < 0.05, **p < 0.01, ***p < 0.001 and ****p < 0.0001. n. s. denotes not significant.

### Author Contributions

R.D. and J.-L.L. designed the study. R.D. and L.-Y.L. performed the experiments. R.D. did data analyses. R.D., L.-Y.L., and J.-L.L. analyzed the results and discussed interpretations. R.D. and J.-L.L. wrote the manuscript.

## Acknowledgments

The plasmid, pAH237, was a kind gift from Professor Tomoyasu Sugiyama at ShanghaiTech University. We thank the Molecular Imaging Core Facility and Molecular and Cell Biology Core Facility at the School of Life Science and Technology, ShanghaiTech University for providing technical support.

## Supplementary information

**Supplementary Table S1.** *S. pombe* strains.

| Fission Yeast Strain | Genotype |
| --- | --- |
| JLL003S | *h- his-D1 ura4-D18 leu1-32 ade6-M216* |
| JLL005S | *h- cts1-YFP (ura4+) ura4-D18 leu1-32 ade6-M216* |
| JLL132R | *h- cts1-H359A-YFP (ura4+) ura4-D18 leu1-32 ade6-M216* |
| JLL127R | *h- cts1-YFP (leu2+) ura4-D18 leu1-32 ade6-M216* |
| JLL128R | *h- cts1-H359A-YFP (leu2+) ura4-D18 leu1-32 ade6-M216* |
| JLL122R | *h- cts1-YFP (ura4+) ura4-D18 leu1-32 ade6-M216/ ade6<sup>+</sup>-htb1-mCherry* |
| JLL123R | *h- cts1-H359A-YFP (ura4+) ura4-D18 leu1-32 ade6-M216/ ade6<sup>+</sup>-htb1-mCherry* |
| JLL125R | *h- cts1-YFP (ura4+) ura4-D18 leu1-32 ade6-M216/ leu<sup>+</sup>-cts1-mCherry* |
| JLL126R | *h- cts1-YFP (ura4+) ura4-D18 leu1-32 ade6-M216/ leu1<sup>+</sup>-cts1-H359A-mCherry* |
| JLL129R | *h- cts1-YFP (ura4+) ura4-D18 leu1-32 ade6-M216/ leu<sup>+</sup>-mCherry* |
| JLL134R | *h- cts1-H359A-YFP (ura4+) ura4-D18 leu1-32 ade6-M216/ ade6<sup>+</sup>-slm9-mCherry* |
| JLL136R | *h- cts1-H359A-YFP (ura4+) ura4-D18 leu1-32 ade6-M216/ ade6<sup>+</sup>-mCherry* |
| JLL135R | *h- cts1-YFP (ura4+) ura4-D18 leu1-32 ade6-M216/ ade6<sup>+</sup>-mCherry* |

**Supplementary Table S2.**
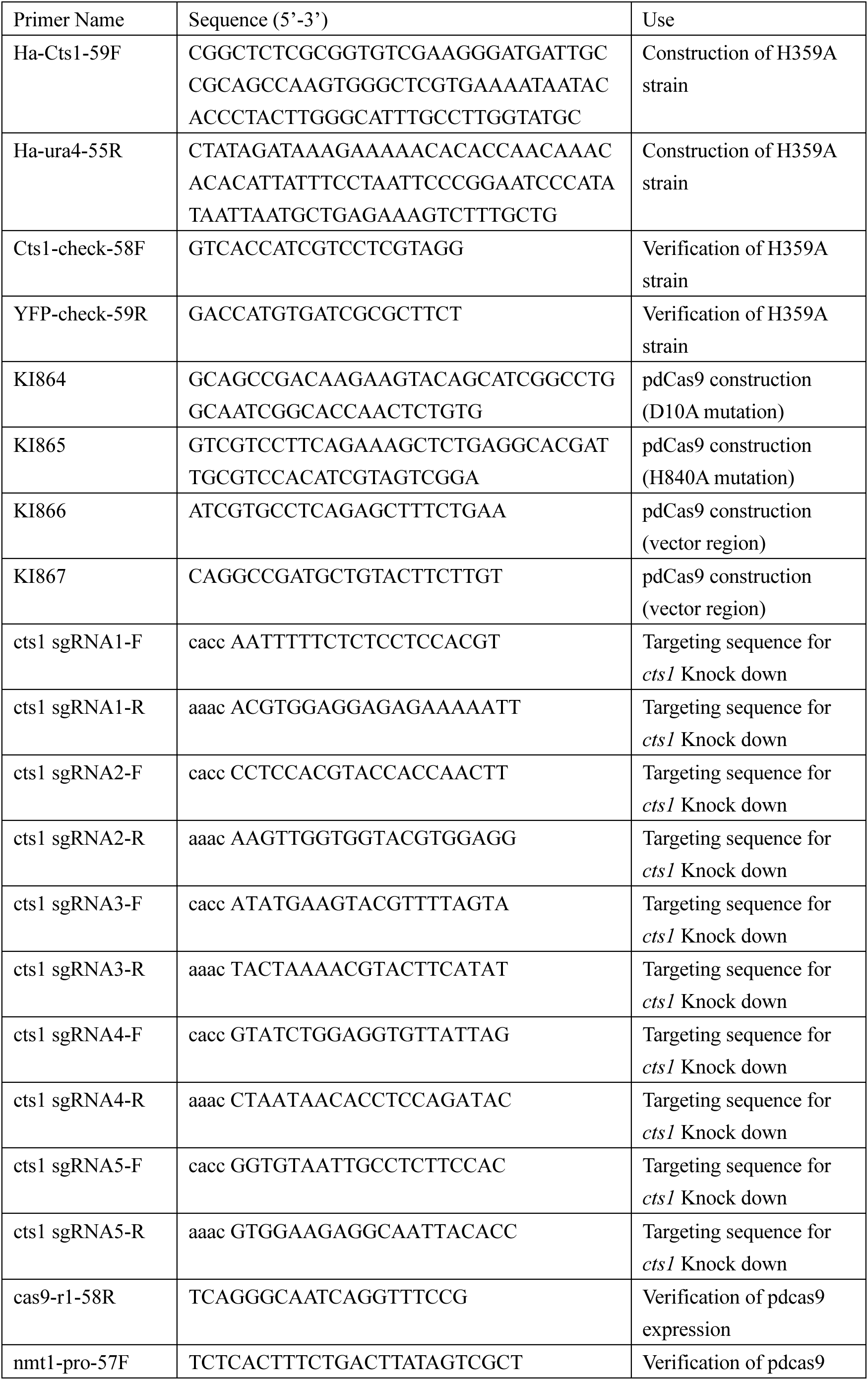
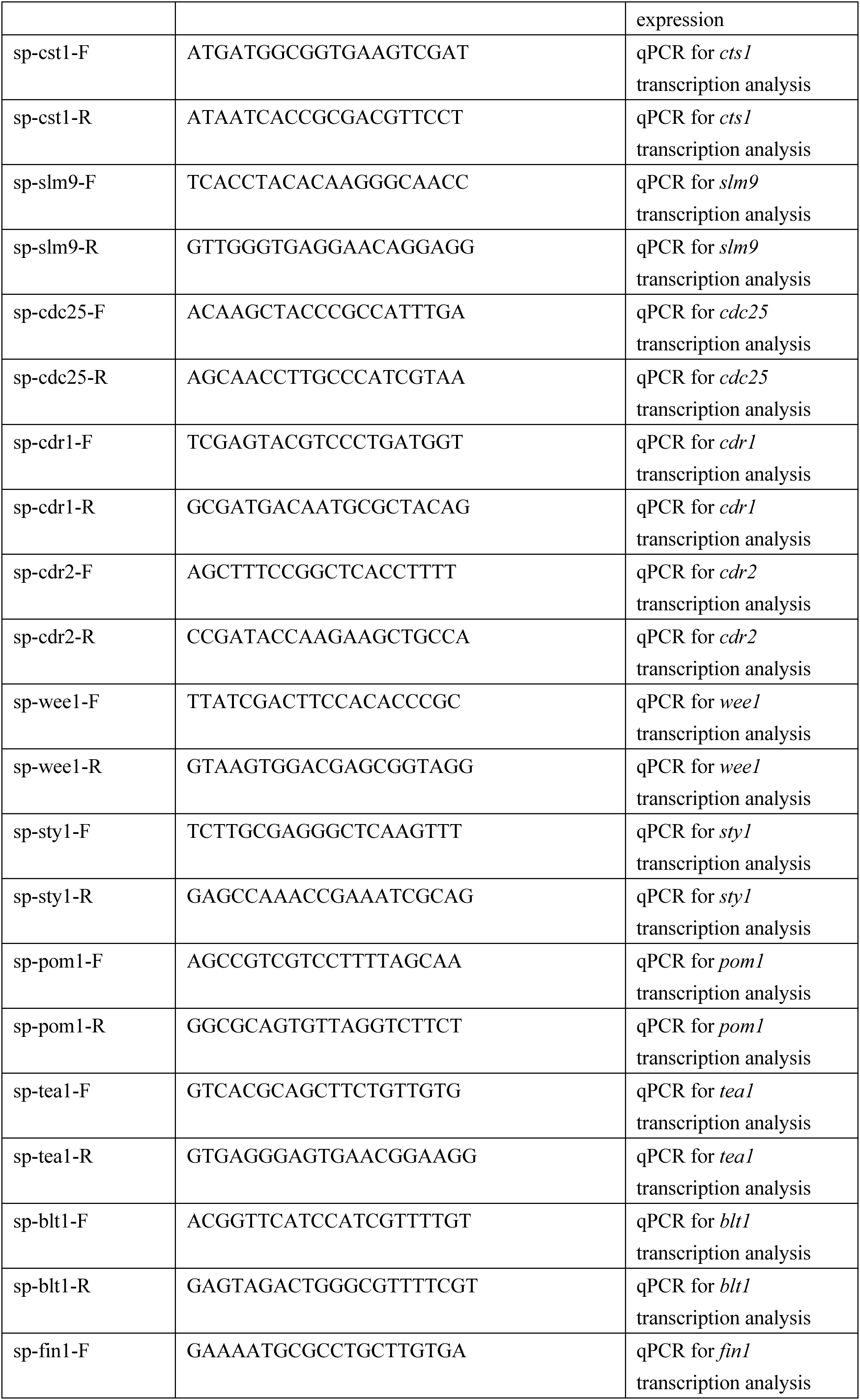
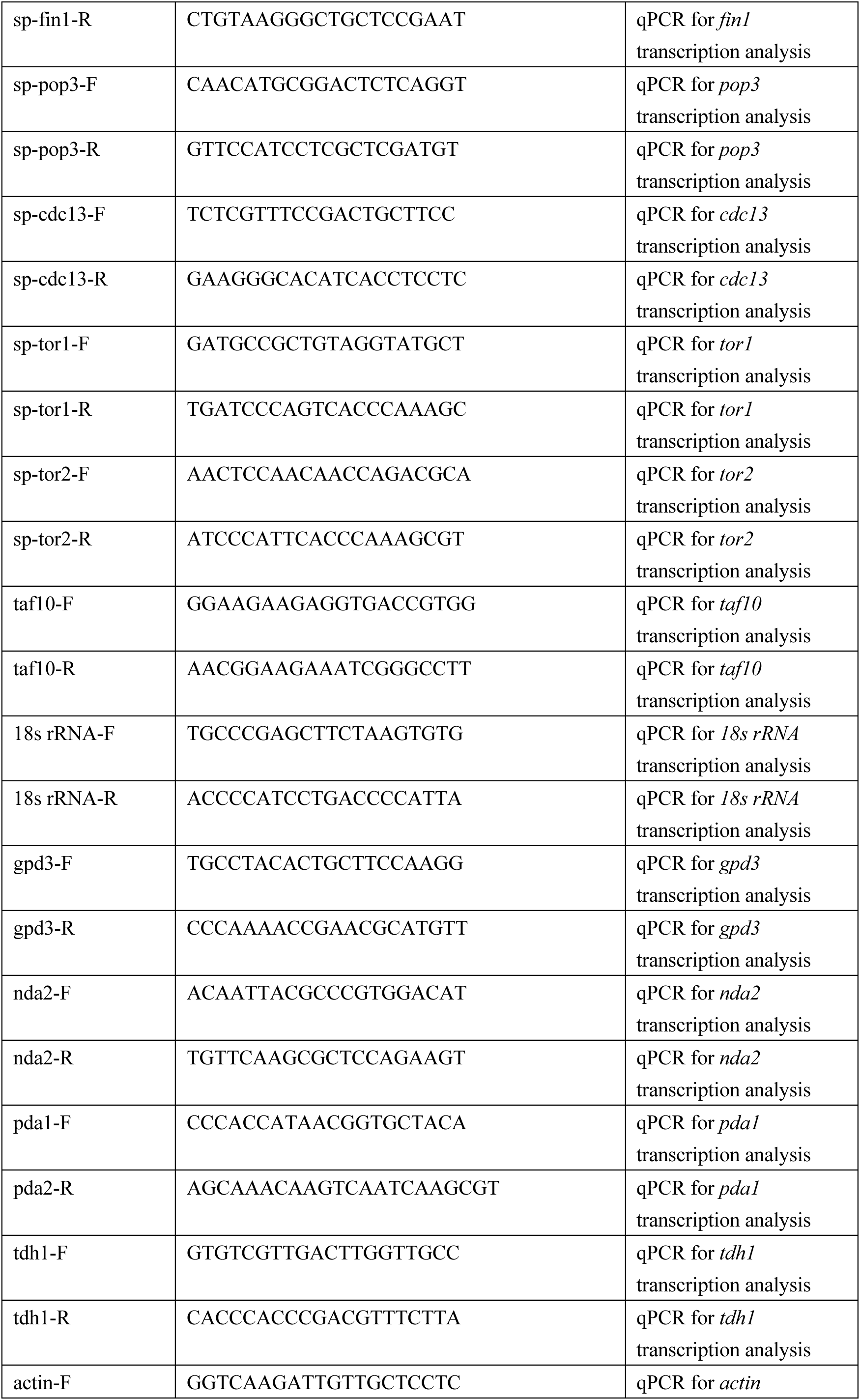
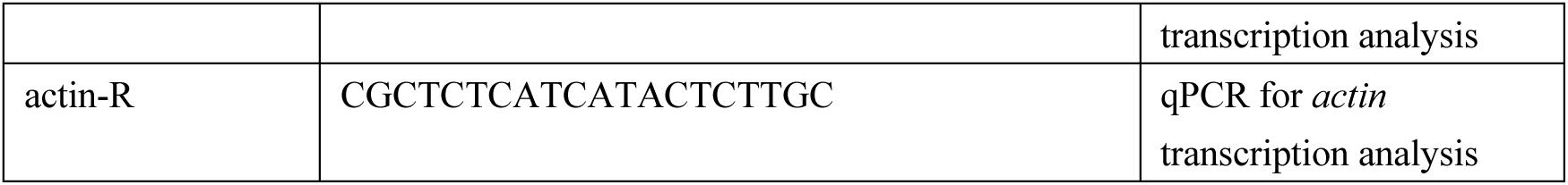
List of primers.

**Supplemental Figure 1.**
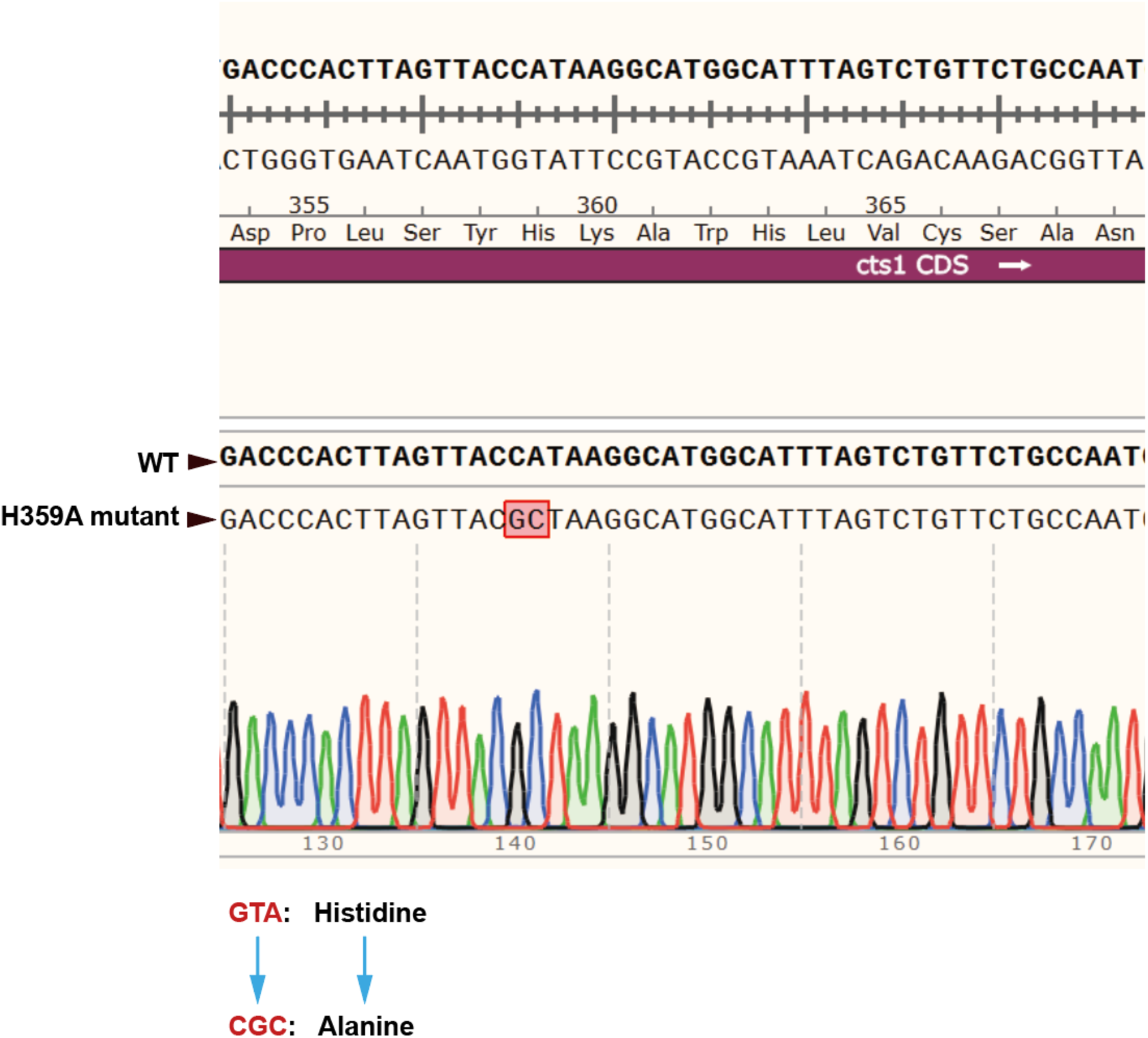
Sequencing analysis. Sequencing analysis of genomic DNA to confirm CTPS H359A point mutation in CTPS^H359A^-YFP strain.

**Supplemental Figure 2.**
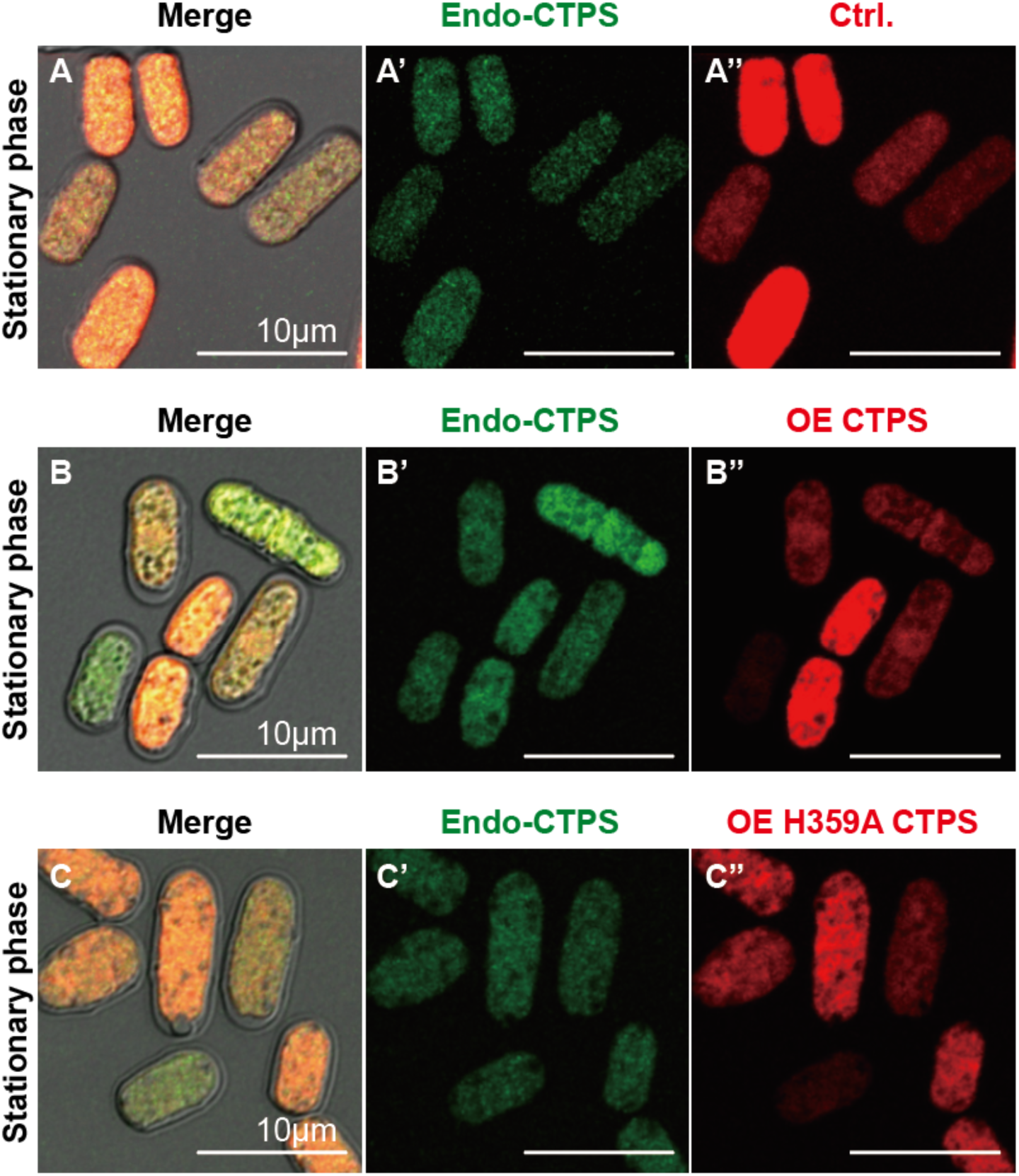
Cytoophidia do not form at stationary phase in various fission yeast strains. **(A-C’’)** Photos of CTPS-YFP strains that overexpression mCherry (A-A’’), CTPS-mCherry (CTPS^WT-OE^, B-B’’), and CTPS^H359A^-mCherry (CTPS^H359A-OE^, C-C’’) at the stationary phase. Cytoophidia did not appear in all strains at the stationary phase. Scale bars, 10 μm.

**Supplemental Figure 3.**
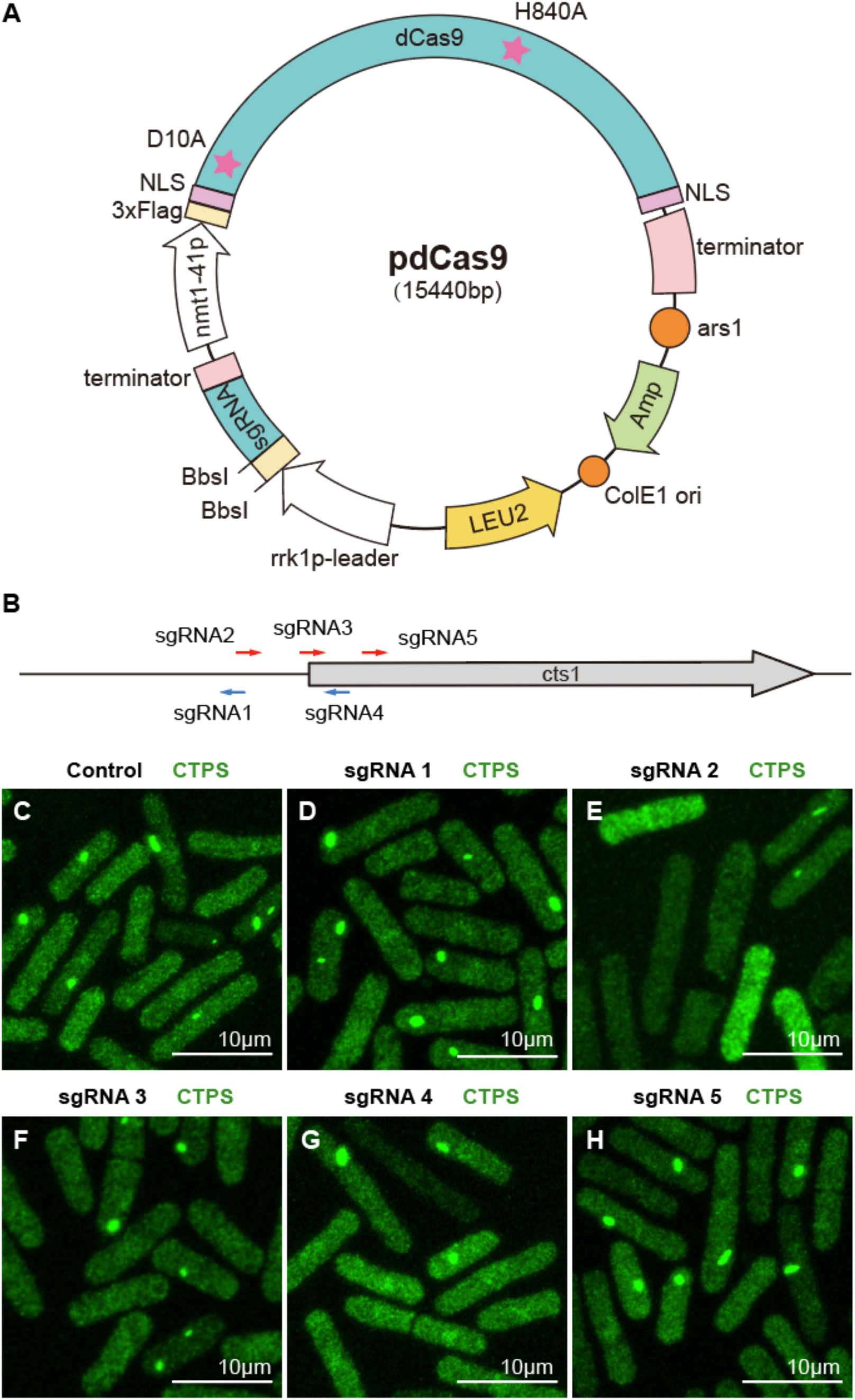
knockdown the CTPS by using the CRISPR-dCas9 system. **(A)** Plasmid map of pdCas9. dCas9, humanized dCas9, originally from *Streptococcus pyogenes*. The pentagram indicated mutations introduced in the dCas9 gene to cause the indicated amino acid substitutions. nmt1-41p, a moderate-strength variant of the nmt1 promoter, the transcription of which is inducible. When the targeting sequence is not replaced, pdCas9 expresses a sgRNA with a nonsense targeting sequence in fission yeast. **(B)** Diagram of the designed targeting sequences with the *cts1* gene. Red and blue arrows indicate targeting sequences of sgRNAs that base-pair with the template and non-template strand of the targeted DNA, respectively. **(C)** Photographs of the CTPS-YFP strains which expressed the pdCas9 plasmid carried nonsense sgRNA (as a control group), sgRNA1, sgRNAs2, sgRAN3, sgRAN4, sgRAN5, respectively, at log phase. Scale bars, 10 μm.

**Supplemental Figure 4.**
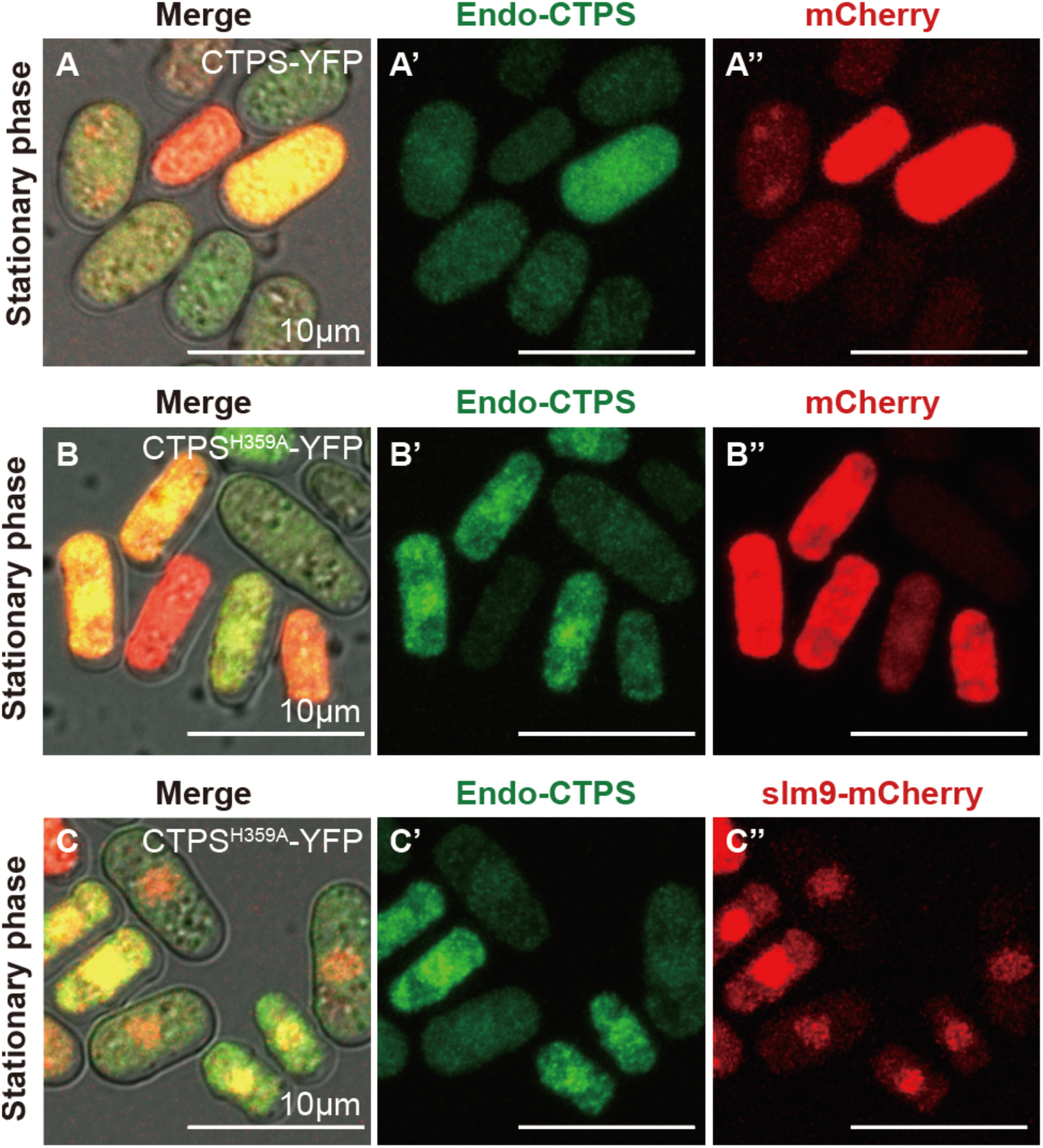
Overexpression of *slm9* in loss-filaments mutant of CTPS strain. **(A-C’’)** Photograph of CTPS-YFP strain that overexpressed only mCherry (named Control.1, A-A’’), CTPS^H359A^-YFP strain that overexpressed only mCherry (named Control.2, B-B’’), and CTPS^H359A^-YFP strain that overexpressed slm9-mCherry, SLM9 localized to the nucleus (named OE-*slm9*, C-C’’) at the stationary phase. Scale bars, 10 μm.

